# Evaluation of Compatibility of 16S rRNA V3V4 and V4 Amplicon Libraries for Clinical Microbiome Profiling

**DOI:** 10.1101/2020.08.18.256818

**Authors:** Po-Yu Liu, Wei-Kai Wu, Chieh-Chang Chen, Suraphan Panyod, Lee-Yan Sheen, Ming-Shiang Wu

**Affiliations:** Department of Internal Medicine, National Taiwan University College of Medicine, Taipei, Taiwan; Department of Internal Medicine, National Taiwan University Hospital Bei-Hu Branch, Taipei, Taiwan; Institute of Food Science and Technology, National Taiwan University, Taipei, Taiwan; Division of Gastroenterology and Hepatology, Department of Internal Medicine, National Taiwan University Hospital, Taipei, Taiwan; Graduate Institute of Clinical Medicine, National Taiwan University College of Medicine, Taipei, Taiwan

## Abstract

Sequencing of the 16S rRNA gene by Illumina next-generation sequencing is broadly used in microbiome studies. Different hypervariable regions of the 16S rRNA gene, V3V4 (amplified with primers 341F–805R) or V4 (V4O; primers 515F–806R), are selected, depending on the targeted resolution. However, in population-based clinical studies, combining V3V4 and V4 data from different studies for a meta-analysis is challenging. Reads generated by short-read (150-bp) high-throughput sequencing platforms do not fully recover the V4 region read-length. Here, we evaluated the compatibility of 16S rRNA V3V4 and V4 amplicons for microbiome profiling. We compared taxonomic compositions obtained by the analysis of V3V4 and V4 amplicons, and V4 fragments trimmed from V3V4 amplicons. We also evaluated an alternative V4 region (V4N; primers 519F–798R) designed for efficient stitching with 150-bp paired-end sequencing. First, we simulated a global investigation of environmental prokaryotes *in silico*. This revealed that V4O primers recovered the highest proportion of fragments (81.7%) and most phyla, including archaea. Empirical sequencing of standard (mock) and human fecal samples revealed biased patterns of each primer that were similar to the ones determined by *in silico* simulation. Further, for human fecal microbiome profiling, the between-sample variance was greater than the systematic bias of each primer. The use of trimmed V4 fragments and single-end amplicons resulted in the same systematic bias. In conclusion, paired-end V4O sequencing yielded the most accurate data for both, simulation and mock community sequencing; the V4O amplicons were compatible with trimmed V4 sequences for microbiome profiling.

**IMPORTANCE:** Next-generation sequencing of the 16S rRNA gene is a commonly used approach for clinical microbiome studies. Different amplicons of the 16S rRNA hypervariable regions are used in different studies, which creates incompatible sequence features when comparing and integrating data among studies by using 16S denoising pipelines. Here we compared the type of data and coverage obtained when different 16S rRNA amplicons were analyzed. *In silico* and empirical analyses of the human fecal microbiome revealed that the V3V4 amplicons are compatible with V4 amplicons after trimming up to the same region. These observations demonstrate that reconciling the compatibility of clinical microbiome data from different studies improve not only the sample size but also the confidence of the hypothesis tested.

## INTRODUCTION

The human microbiome affects human health in numerous ways (1-7). Notably, the gut microbiome mediates host metabolic and physiological status, such as digestion, immune response, neuron transmission, and circulation (8). Clinical research of the microbiome is rapidly advancing, propelled by sequencing of the 16S rRNA gene (abbreviated “16S”) using the next-generation sequencing (NGS) technology. Although long-read sequencing technologies have matured in recent years, the well-developed analysis pipelines for massive microbiome taxonomic profiling (e.g., QIIME2) are mainly based on the Illumina sequencer systems (9).

Accurate evaluation of the microbiota heavily depends on the primers used (10, 11). Further, lower-level taxonomic resolution bias can arise when non-representative regions are amplified (12). The 16S V3V4 (primers 341F–805R) and V4 (primers 515F–806R) hypervariable regions are most frequently used for human microbiome profiling (6, 13). When an amplicon library is prepared using Nextera XT two-step polymerase chain reaction (PCR), the expected insert sizes from these regions are 465 bp and 291 bp, respectively. (14, 15). The V3V4 and V4 amplicons are fully recovered on the Illumina MiSeq platform, which can be used to sequence up to 600 nucleotides from both ends of an amplicon [(300 bp)×2].

When the National Institute of Health Human Microbiome Project (HMP) started in 2008, investigations of the human microbiota using Roche/454 pyrosequencing focused on the 16S V3V5 region (primers 357F–926R) (16). However, the V3V4 region became the mainstream amplicon target in microbiota studies since Illumina released a recommended library preparation protocol for sequencing on the MiSeq platform (15). Although the outputs of both sequencing approaches were comparable (17), the Illumina platform generated much more reads than pyrosequencing. After HMP, in 2010, the Earth Microbiome Project (EMP) was initiated, as a global investigation of environmental and host-associated microbiomes (18, 19). The 16S V4 region (primers 515F–806R) was amplified for sequencing in the EMP projects (18), including human-associated microbiome studies (20) and the American Gut Project (21). This allowed for a more representative prokaryotic profiling since the V4 universal primer pair effectively captures both bacterial and archaeal 16S sequences. However, the EMP V4 libraries were constructed using custom sequencing primers according to Caporaso et al. (22), and generated by 150-bp paired-end (PE) sequencing with additional 14 cycles for barcode tagging on MiSeq or HiSeq. The amplicon insert size of the custom V4 library is expected to be 252 bp, while the two-step PCR method generates 291 bp from the same targeted region (14). This means that the Nextera XT two-step PCR method is only suitable for platforms with an over 150-bp PE sequencing capacity (250-bp PE and 300-bp PE). Nonetheless, the V4 region is analyzed regardless of which protocol is followed. Data from different protocols are supposed to be ideally integrated amplicon sequence variants (ASVs) for meta-analysis microbiota profiling by using denoising algorithms (23) [e.g., DADA2 (24), Deblur (25), and UNOISE3 (26, 27)] in the QIIME2 pipeline (28).

The sequencing cost per megabase has exceeded Moore’s Law, which describes a trend in doubling of computing power that conceives the improvement of DNA sequencing capacity yearly and even over the exponential trend (29). The Illumina sequencing platforms generate between several tens to hundreds of millions of reads, enabling deep profiling of a large number of samples during a single PE run at a fraction of the cost of a study. The output of more than 100,000 reads per sample is suggested and sufficient for microbiota investigations (15). Reduced sequencing runs are preferred in large sample-size studies, especially clinical cohort studies, to avoid batch effects. Higher throughput sequencers, such as NextSeq and HiSeq, which are common in academic core laboratories, generate reads with a low batch effect and at a low cost per sample with a single run (30). However, the confidence for stitching both ends to recover the full V4 region after quality trimming of thus generated PE sequences [(150 bp)×2, the maximum read length of NextSeq] is low. Hence, it is difficult for the higher throughput platforms to meet the demands for cost–benefit and data yields.

In response to the changes of amplicon sequencing methods in the clinical gut microbiota research and considering the cost–benefit of sequencing, we here compared several pairs of sequencing and analysis approaches. We first conducted an *in silico* PCR simulation of a global investigation of environmental prokaryotes by capturing the 16S V3V4 and V4 regions, as well as the V4 fragments trimmed from V3V4 from the SILVA 132 ribosomal RNA NR 99 database (DB). We tested primer and analysis bias by analyzing a mock microbial community. We then sequenced human fecal samples and analyzed the data using the QIIME2 pipeline with the DADA2 plugin, which is currently the most accurate sample inference method (denoising). We then evaluated the compatibility (including the taxonomic abundance consistency and coverage) of the amplicons of different 16S hypervariable regions. The analysis revealed primer-and analysis method-associated bias. Based on the findings, we propose optimized analytical options for clinical population-based and meta-analysis studies.

## RESULTS

### Cost evaluation of 16S amplicon sequencing using the Illumina MiSeq and NextSeq platforms

NGS sequencers are versatile platforms for sequencing-based studies. The cost– benefit ratio (output quantity and quality) of sequencing has to be considered when choosing a suitable platform. Focusing on the 16S amplicon sequencing, we compared the cost effectiveness of the Illumina mid to mid–high throughput platforms (MiSeq and NextSeq platforms), which the majority of academic core laboratories are commonly equipped with (Table 1).

**TABLE 1.** Summary of the estimated 16S amplicon sequencing throughputs and costs for MiSeq and NextSeq platforms.

|  | MiSeq | NextSeq, mid-output | NextSeq, high-output |
| --- | --- | --- | --- |
| Flow cell configuration | 600 cycles (300-bp PE) | 300 cycles (150-bp PE) | 300 cycles (150-bp PE) |
| Throughput | 13.2–15 Gb (44–50M reads) | 32.5–39 Gb (216–260M reads) | 100–120 Gb (660–800M reads) |
| Data quality<sup>a</sup> | >70%, >Q30 | >75%, >Q30 | >75%, >Q30 |
| Targeted 16S hypervariable regions | V3V4, V4 | V4 | V4 |
| No. samples | 175 | 384 | 384 |
| per run per 100K PE reads<br>(Nextera XT 2-Step PCR)<sup>b</sup> | (out of the possible 384 samples) | (39.4% read output of a single run) | (12.8% read output of a single run) |
| Maximum no. samples | 175 |  | 2168 |
| per run per 100K PE reads<br>[Caporaso et al. (36), protocol, 1-Step PCR]<sup>c</sup> | (out of possible 2168 samples) | 975<br>(out of possible 2168 samples) | (all possible index combinations, 72% read output of a single run) |
| Cost of sequencing<br>per 100K-read output per sample<sup>d</sup> | US\$15.2 | US\$3.4 | US\$2.8 |

<sup>a</sup>Data quality scores (average % of bases over the entire run), as reported by Illumina documents (15).
<sup>b</sup>Quality read outputs were determined as maximum Q30 proportion, 291-bp V4 insert size, and less than 10-bp overlap of 150-bp PE reads. PE: paired-end.
<sup>c</sup>V4 insert size of 252 bp.
<sup>d</sup>The costs (in US dollars) were determined based on the prices cited by the Medical Microbiota Center of the First Core Laboratory, National Taiwan University College of Medicine (Taipei, Taiwan).

The maximal flow cell configurations of MiSeq and NextSeq are 600 (300-bp PE) and 300 (150-bp PE) cycles, respectively. Based on the Illumina Nextera XT library preparation method for 16S amplicons, the V3V4 and V4 regions are fully recovered using the MiSeq 600-cycle reagent kits; on the other hand, only the V4 region can be in “theory” recovered using the NextSeq 300-cycle reagent kits. Fixing the output at 100K PE reads per sample, the MiSeq can process 175 samples per run (96 samples per run as the regular configuration), using the default library preparation method (Nextera XT). The mid-output and high-output flow cells of the NextSeq can process up to 384 samples per run (all index pairs), accounting for 39.4% and 12.8% reads of a single run, respectively. The 16S amplicon sequencing costs in US dollars are $15.2 per 100K reads per sample for the MiSeq, decreasing to $3.4 and $2.8 for the NextSeq mid-output and high-output platforms, accordingly.

Since the Illumina NGS base quality decreases toward the 3’-end over the read (Fig. 1A), the overlapping area of the PE sequences should be sufficiently long to allow the assembly by high-quality base pairs (Fig. 1B). However, when the V4 amplicon sequencing is performed via short read-lengths (e.g., 150-bp PE), gaps exist between the two amplicon ends after quality trimming (Fig. 1C). Caporaso et al. (29) modified the V4 library preparation method to make it suitable for use with 150-bp PE sequencing. The modified method generates 252-bp amplicons instead of 291-bp amplicons obtained with the Nextera two-step PCR approach. However, although the modified method increases both, the V4 amplicon assembly efficacy and sequencing capacity (up to 975 and 2168 samples per run on the NextSeq mid-output and high-output platforms), it requires manual alteration of the sequencing software configuration. The above comparisons revealed that it is necessary to optimize sequencing configurations (i.e., library construction by amplifying suitable hypervariable region and sequencing with long enough configuration) for improving the cost-benefit of 16S amplicon sequencing. We, therefore, further proceeded to test if the shorter (V4) amplicons harbor equivalent or better taxonomic profiling capacities compared to the longer (V3V4) amplicons.

**FIG 1.**
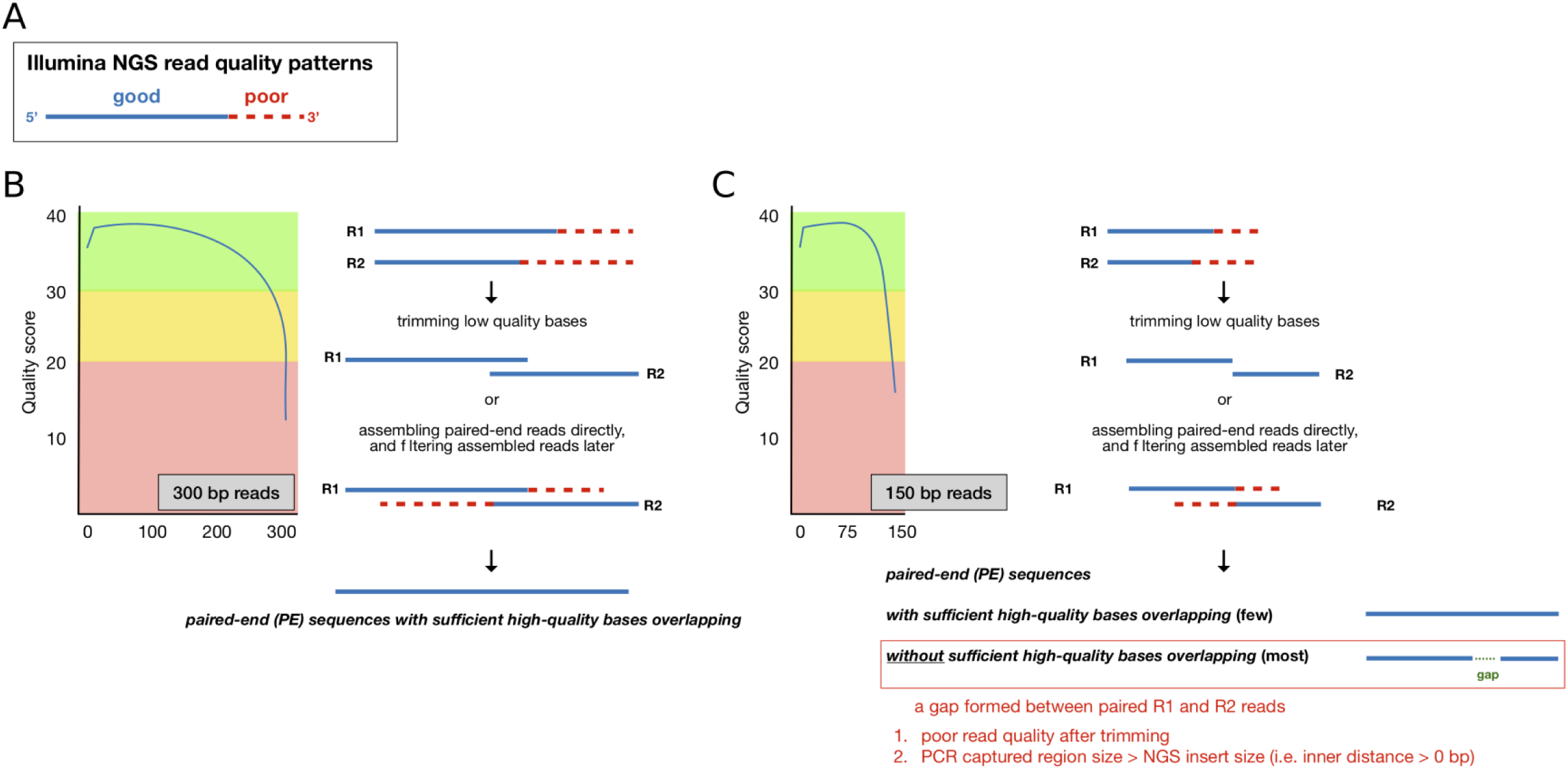
NGS read-quality pattern diagram. (A) Illumina read-quality patterns: from the 5’-end to 3’-end, the quality per base decreases and is usually trimmed (red dashed line) before assembly. (B) The 300-bp paired-end (PE) read-quality distribution and amplicon stitching procedures. The 300-bp PE reads with sufficiently long overlapping regions are stitched together. (C) The 150-bp PE read-quality distribution and amplicon stitching procedures. Only a few read-pairs pass the quality filtering and assembly at both ends. Most 150-bp PE reads either fail to pass the quality control or fail to recover amplicon inserts that are too long.

### Comparison of taxonomic profiling capacities of the 16S V3V4 and V4 regions by an *in silico* PCR simulation

To evaluate the primer efficacies of 16S V3V4 (primers 341F–805R) and V4 primer pairs (including V4 original, V4O, primers 515F–806R; alternative V4, V4N, primers 519F–798R; tV4O, V4O region trimmed from V3V4 fragment; and tV4N, V4N region trimmed from V3V4 fragment), we simulated PCR capture of the targeted fragments from the DB [SILVA 16S gene database (NR 132 99%)](31, 32). The simulation encompassed an investigation of the primer-dependency of detected bacterial and archaeal profiles because the *in silico* PCR captured all targeted sequences from the global environment.

The DB contains 369,953 representative 16S sequences (Table 2). Our targeted approaches extracted 59.2% to 81.7% of sequences from the DB. Although the V3V4 primers captured the longest fragments, they extracted 77.3% of all DB sequences, while the V4O primers extracted 81.7% of all sequences. The V4N-captured sequences covered most of the V4O region but were shorter by approximately 11 bp. This reduced the capture rate of V4N to 66.2%. The tV4 primers recovered 72.5% (tV4O) and 59.2% (tV4N) of sequences from the DB, and 93.8% (tV4O) and 76.5% (tV4N) of the V3V4 sequences.

**TABLE 2.**
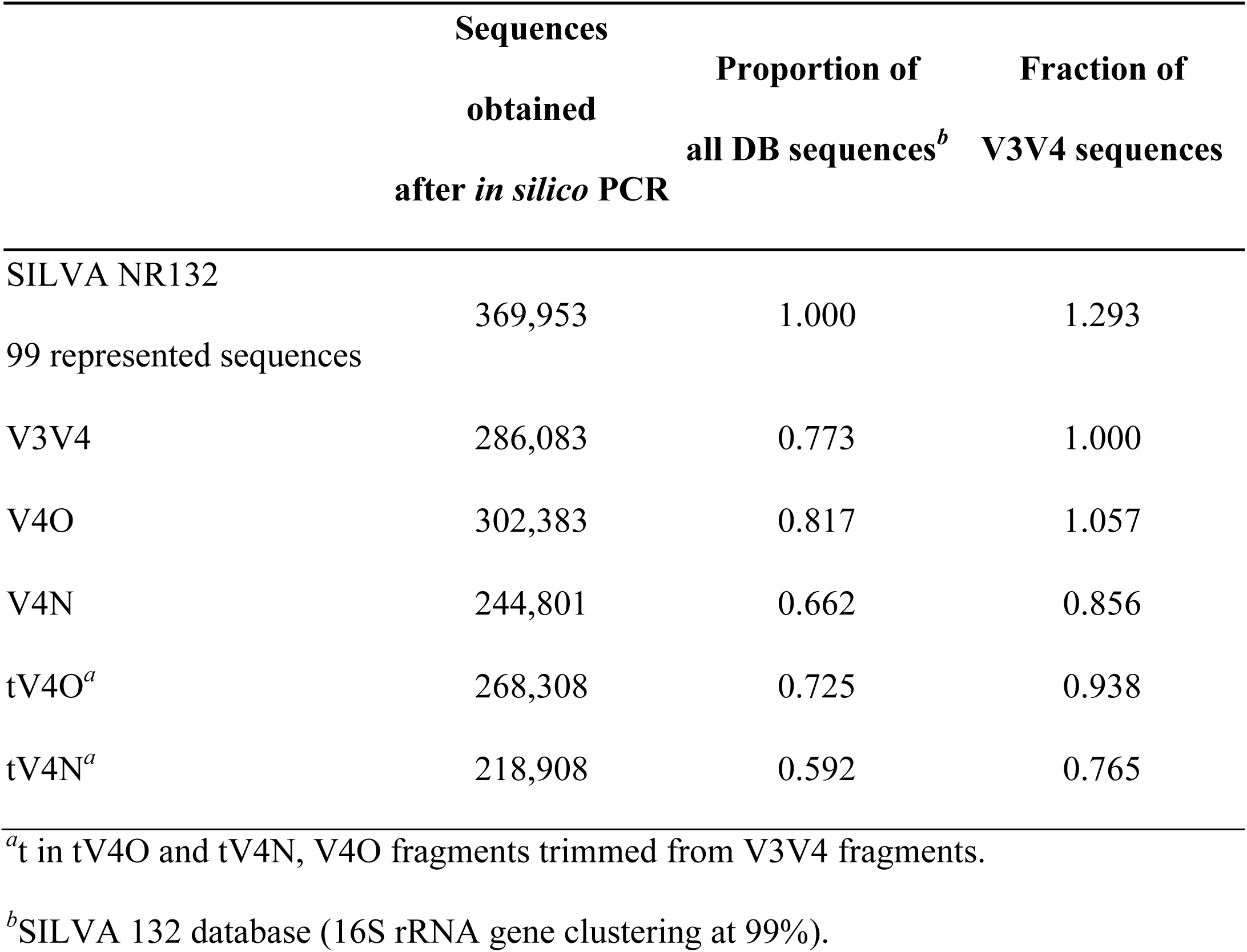
Coverage rates of *in silico* PCR simulation with the V3V4 and V4 primers

Based on the assigned taxonomy analysis, all primers captured similar proportions of the major phyla in *in silico* PCR (Fig. 2A). However, the overall differences from the DB composition ranged from 6.44% to 36.69% of the phyla proportions. The V4O resulted in the least differences (6.44%) from the DB, followed by V3V4 (15.23%) and V4N (30.0%). The trimmed V4 approaches led to datasets that differed by 16.9% (tV4O) and 36.69% (tV4N) from the DB. Both V4O and V4N approaches captured the maximum archaea (5.26% and 3.40%, respectively; Table S1), while the V3V4 approach captured the fewest archaea (0.02%). The two trimmed V4 approaches, designed based on V3V4, did not efficiently capture the archaea (0.02% for tV4O and 0.005% for tV4N).

**FIG 2.**
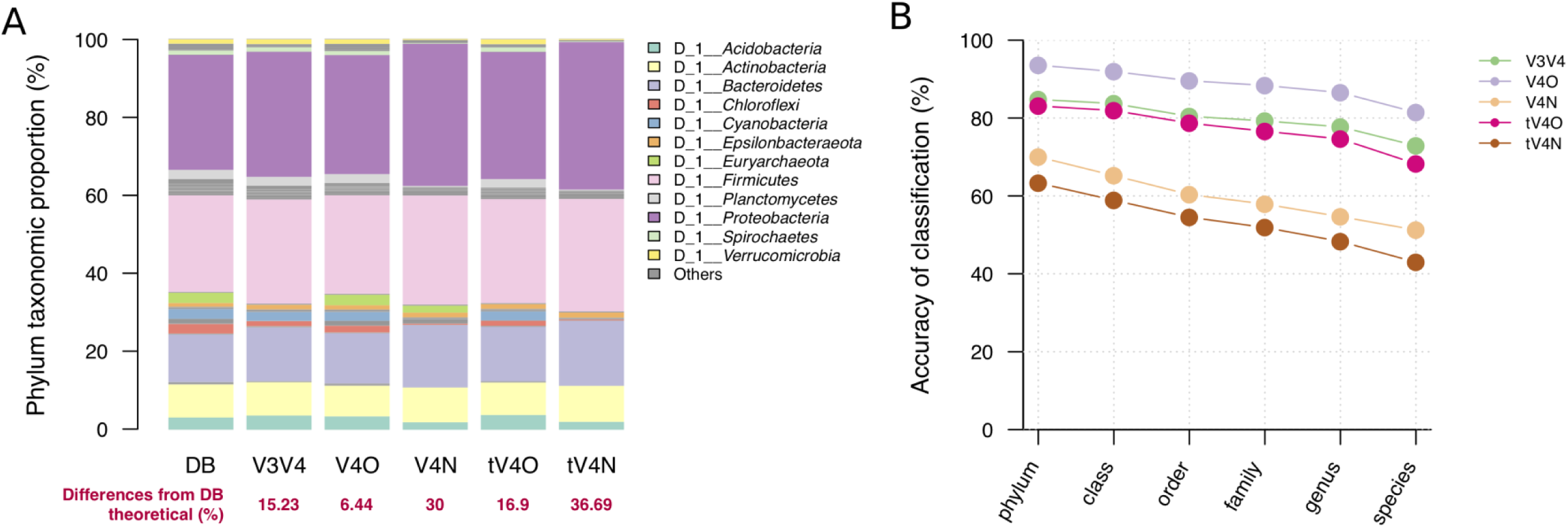
Taxonomic profiling via *in silico* PCR simulation. (A) Phylum composition of the SILVA 16S database and extracted phylum composition for the five tested conditions of *in silico* PCR. (B) Phylum to species taxonomic accuracies (against taxonomic assignment in the SILVA 16S database) for the five conditions of *in silico* PCR. The five conditions for *in silico* PCR were as follows:V3V4 (341F–805R), V4O (515F–806R), V4N (519F–798R), tV4O (trimmed V4O from V3V4), and tV4N (trimmed V4N fromV3V4).

We then compared the accuracy of the classification at each taxonomic level by the different primer approaches (Fig. 2B). Over 90% of V4O-generated sequences were assigned correct taxonomy, followed by the V3V4 and tV4O sequences (approximately 80% accuracy at each level). However, at most 70% and 63% of the V4N and tV4N sequences, respectively, were assigned the correct taxonomy (summarized in Table 3). This indicates that the V4O primer would yield the best taxonomic profiles in a global microbial investigation.

**TABLE 3.**
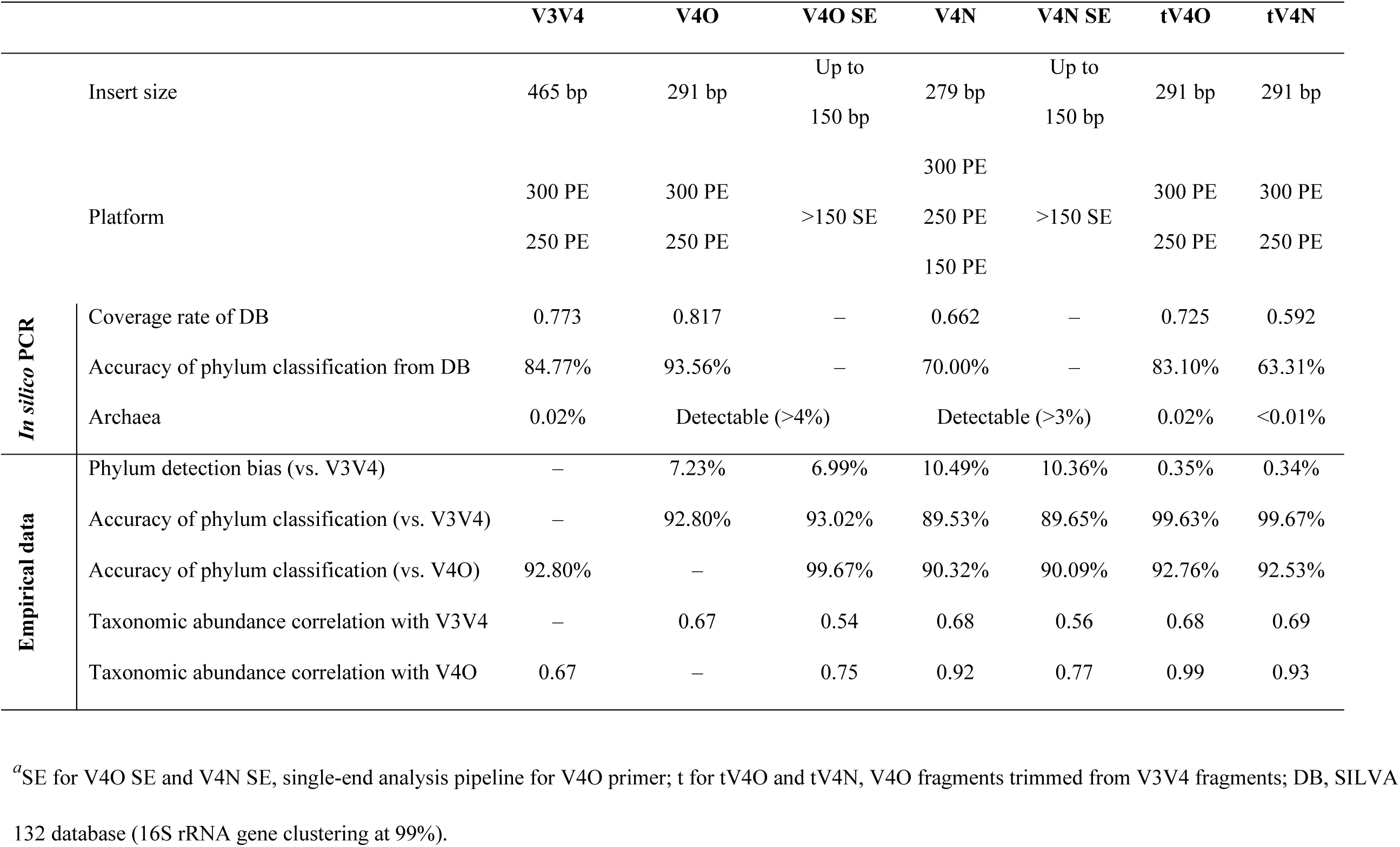
Summary of sequencing efficacies (*in silico* and empirical data), considering the coverage rate, taxonomic identification rate, and taxonomic abundance, and depending on the primers and analysis approaches^*a*^

### Mock community profiling by using seven different 16S amplicon analytic approaches

To empirically evaluate the compatibility of V3V4 and V4 primers, we sequenced and analyzed a mock microbial community. The mock community represented eightpspecies of bacteria and two species of yeasts (see Materials and Methods) and is an artificial synthesized microbial community that serves as a quantitative standard. We used seven analytical approaches, all in conjunction with the QIIME2 DADA2 denoising pipeline (9, 24)(Fig. 3), i.e., we analyzed PE V3V4 amplicons (V3V4), PE V4O amplicons (V4O), PE V4N amplicons (V4N), single-end V4O amplicons (V4OSE), single-end V4N amplicons (V4NSE), and V4 amplicons trimmed from V3V4 (trimmed V4O, tV4O; and trimmed V4N, tV4N). We also sequenced human fecal samples from 10 volunteers using the same protocols utilized to evaluate the primer compatibilities in real targeted samples.

**FIG 3.**
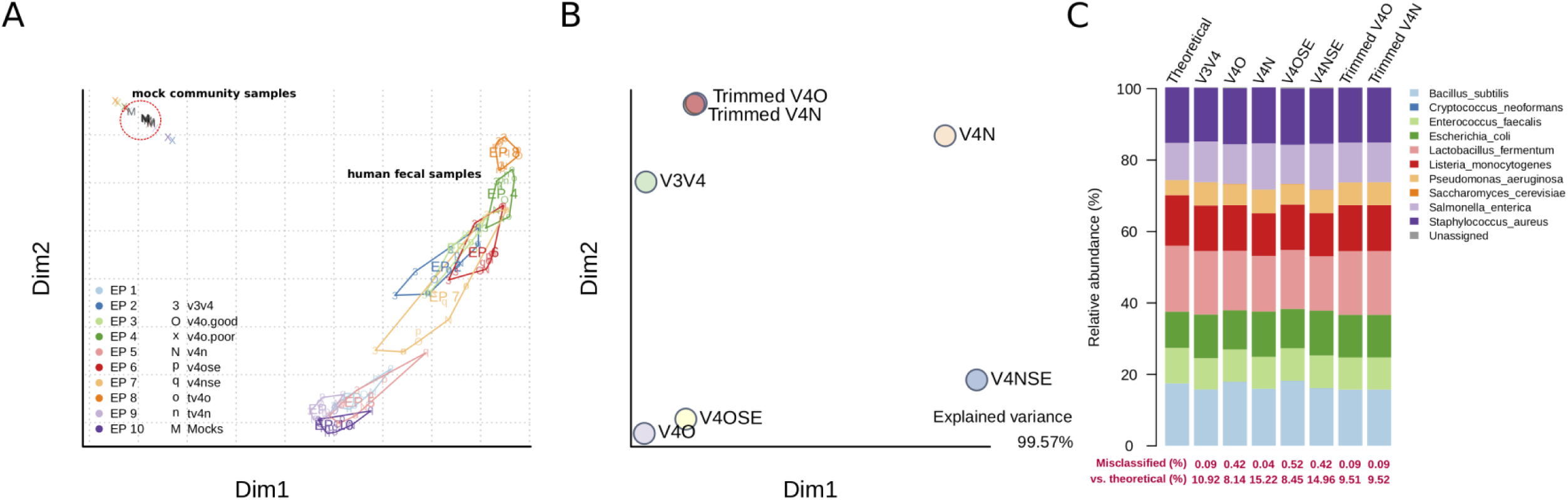
Compatibility evaluation for three primer-amplification and seven analytical approaches, using a mock microbial community sample. (A) Beta-diversity ordination (Bray–Curtis dissimilarity) of the human fecal microbiome (EP1–EP10; colored symbols) and mock community (M; black symbols) sequencing samples. Notation: 3, V3V4 amplicon library; O, successful assembly of V4O amplicon library; x, failed assembly of V4O amplicon library; N, V4N amplicon library; p, V4OSE, single-end analysis using V4O amplicon library; q, V4NSE, single-end analysis using V4N amplicon library; o, tV4O, trimmed V4O amplicon from V3V4 library; n, tV4N, trimmed V4N amplicon from V3V4 library. The mock community sample analyzed by seven analytical approaches clustered together and away from data for 10 individual human fecal microbiomes. (B) Beta-diversity ordination of the seven analytical approaches used for the analysis of mock community data. (C) The relative abundance of bacteria in the mock community, comparing the theoretical composition with that obtained by seven analytical approaches.

First, we compared the data for mock community samples (simple sample composition) with human fecal samples (complex sample composition) using the seven analytic approaches (Fig. 3A). The beta-diversity (measured by Bray–Curtis dissimilarity) ordination plot revealed high homogeneity in mock community quantification of the seven approaches. The Bray–Curtis dissimilarity is a quantitative measurement. Accordingly, a zoomed-in view of the beta-diversity ordination for the mock community revealed that the PCR conditions (primers used) reflected the community quantitative composition (Fig. 3B). Specifically, the trimmed V4 data points were close to the V3V4 data point; the PE and SE V4O data points were clustered together; and the PE and SE V4N data points were proximal on the first plot axis.

The relative taxonomic abundances in the theoretical sample, as determined by the seven analytical approaches, were not significantly different (G-test; mean of G=1.31, P=0.99; Fig. 3C). The rates of misclassification (unassigned or assigned to yeasts) were less than 1% in all analyses. The differences in the determined relative abundance and the theoretical composition ranged from 8.14% (V4O) to 15.22% (V4N). Sample composition determined by the V3V4 approach (and its derived V4 approaches) differed by approximately 10% from the theoretical composition. This indicates that the different 16S regions and analytic approaches for profiling a simple composition microbiota, such as a synthesized mock community, are able to present similar quantitative and qualitative results.

### Determination of sample heterogeneity and variation of taxa composition after 16S amplification with different primers

We next analyzed human fecal samples to evaluate the complementarity of different 16S amplification analytical approaches. The seven analytical approaches used were the same as those for the mock community analysis. We examined the beta diversity, as determined by the seven approaches (Fig. 4). The beta-diversity profiles for fecal microbiota data points for the seven approaches reflected the 10 individual sample sources (Fig. 4A). We tested the sample and analytical approach heterogeneity by the analysis of similarities (ANOSIM). The between-sample variance was greater than within-sample variance (R=0.987, P=0.001; Fig. 4A and Fig. 4B); on the other hand, the between-analytical approach variance was not significantly greater than the within-analytical approach variance (R=0.016, P=0.22; Fig. 4C and Fig. 4D). In other words, the determined sample variation was relatively constant, regardless of the primers or analytical approaches used.

**FIG 4.**
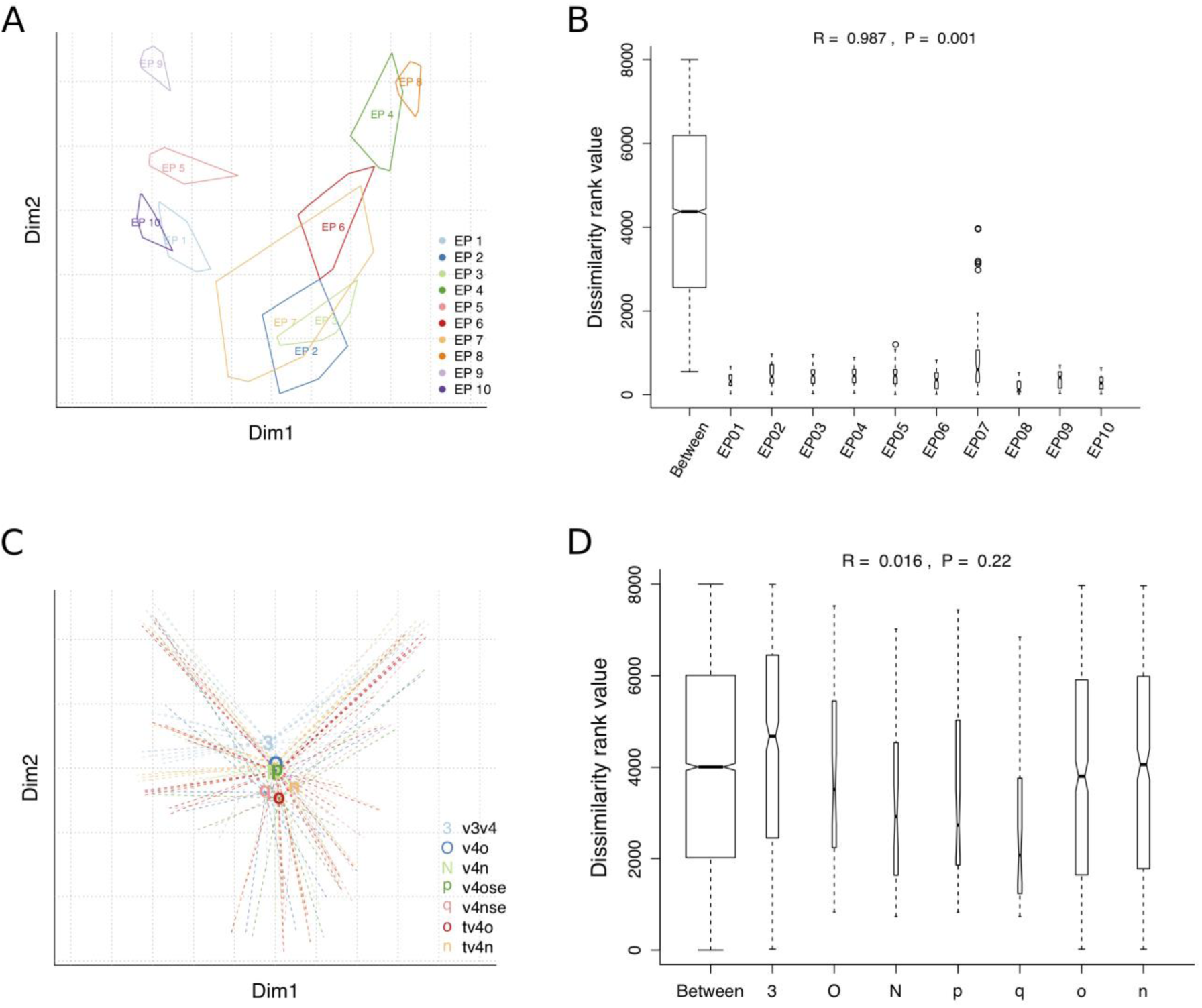
Compatibility evaluation of three primer-amplification and seven analytical approaches by testing the heterogeneity of 10 human fecal microbiomes. (A) Beta-diversity ordination (Bray–Curtis dissimilarity) of 10 human fecal microbiome samples (colored-border polygons, EP1–EP10). (B) Nested analysis of similarity (ANOSIM) of individual testing approaches, nested by the analytical approach; R=0.987, P=0.001. (C) Beta-diversity ordination of 10 human fecal microbiome samples (radial lines; the analytical approaches are labeled in the center). (D) Nested ANOSIM of the analytical approaches nested by individuals; R=0.016, P=0.22.Abbreviations are as in Fig. 3.

We next analyzed three alpha-diversity indices (namely, richness, Shannon entropy, and Simpson index of diversity), as determined by using the seven analytical approaches (Fig. S1). The three indices were not statistically significantly different when the different approaches were used (P>0.05). However, the differences in the richness index were marginally significant (P=0.08) for the seven approaches; this trend was attributed to reduced taxon numbers in samples analyzed using the trimmed V4O and trimmed V4N approaches (Fig. S1A).

### Consistency of taxonomic abundances determined by different 16S amplicon analytical approaches

PCR artifacts (over-amplified amplicons and chimeras) interfere with taxonomic quantification and taxonomic structure profiling of microbiome samples. The artifacts are sequences that are amplified in a biased manner during PCR. No universal primer exists for a fully unbiased amplification. In the current study, we used the DADA2 denoising pipeline to reduce the confounding effect of PCR artifacts. We profiled the higher-level taxonomy compositions (the relative phylum abundance) by using stacked barplots (Fig. 5A). The proportion of each phylum was different for different analytical approach used, but the difference was not statistically significant (G-test; mean of G=0.81, P=1). However, the V4N-PE analytical approach under-detected the phylum *Verrucomicrobia* (relative abundance 0.02% vs. 1% of other approaches).

**FIG 5.**
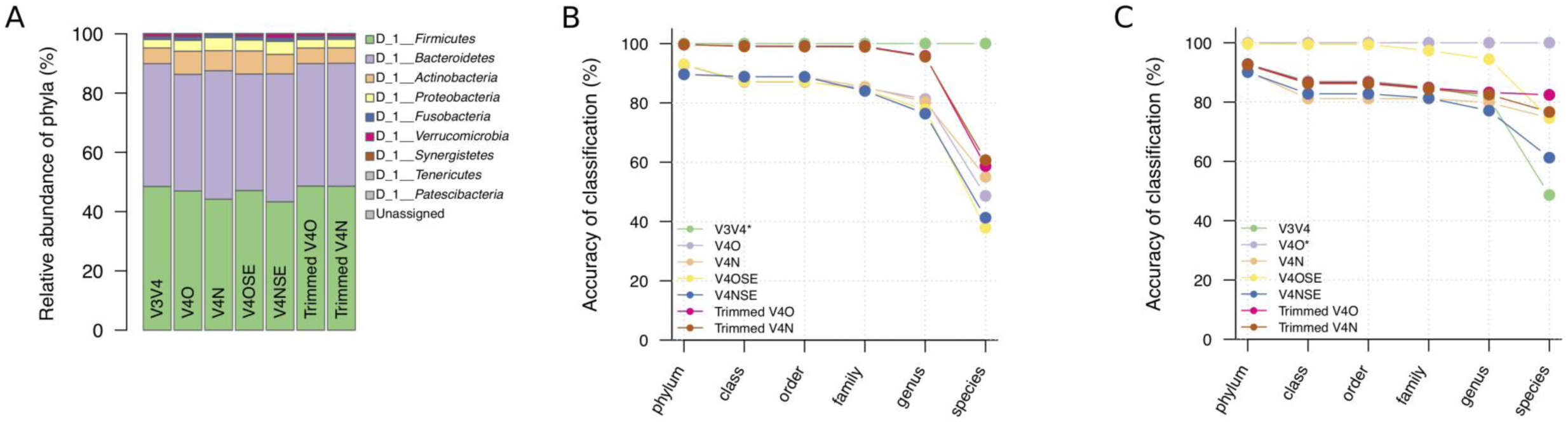
Phylum compositions and taxonomic accuracy of the human fecal microbiomes analyzed by seven analytical approaches. (A) Phylum compositions of the human fecal microbiome (10 individuals average per stacked bar) analyzed by seven analytical approaches. (B) Phylum to species taxonomic accuracies based on the V3V4 taxonomy assignment. (C) Phylum to species taxonomic accuracies based on the V4O taxonomy assignment. Abbreviations are as in Fig. 3.

Because no theoretical reference exists for the human fecal microbiota composition, we used the V3V4 approach (the longest sequence region; Fig. 5B) and the V4O approach (the most accurate method based on *in silico* simulation and mock community experiments; Fig. 5C) as benchmarks to evaluate the classification accuracy. The taxonomy data obtained by V3V4-derived analyses (trimmed V4O and trimmed V4N) were the closest to those of the V3V4 approach above the genus level (Fig. 5B). Compared with the V3V4 data, the accuracy of V4O and V4N (PE and SE) methods above the genus level was 78.9% to 91.3%. On the other hand, compared with the V4O data, the taxonomy determined by the V4OSE method was the closest to that of the V4O (PE) method (Fig. 5C). The other methods reached an 83.1% to 93.0% average relative accuracy above the genus level. The rank abundance analysis at the family (Fig. S2A) and genus levels (Fig. S2B) revealed that the taxon abundance was consistent with the V4O-based classification accuracy.

We then performed pairwise analysis of taxonomic abundance correlations between the seven analytical approaches (Fig. 6). We plotted the abundance of each taxonomy-assigned ASV using pairwise-correlation scatter plots (Fig. 6, lower left panels) and noted the abundance correlation coefficients (Fig. 6, upper right panels). The correlation coefficients between the V3V4 data and other methods ranged from 0.54 to 0.69. The correlation coefficients between the V4 method data, regardless of whether these were PE, SE, or trimmed methods, and the V4O and V4N data were high (0.92 to 0.94). In addition, for the V4 PE data, we observed the best correlation between the corresponding trimmed V4 methods (0.99 for V4O and tV4O; and 0.98 for V4N and tV4N) (summarized in Table 3). A high consistency among V4 approaches presented by integrating both taxonomic assignment and quantification, and the V3V4 amplicons would be compatible with V4 by trimming up to the same hypervariable region.

**FIG 6.**
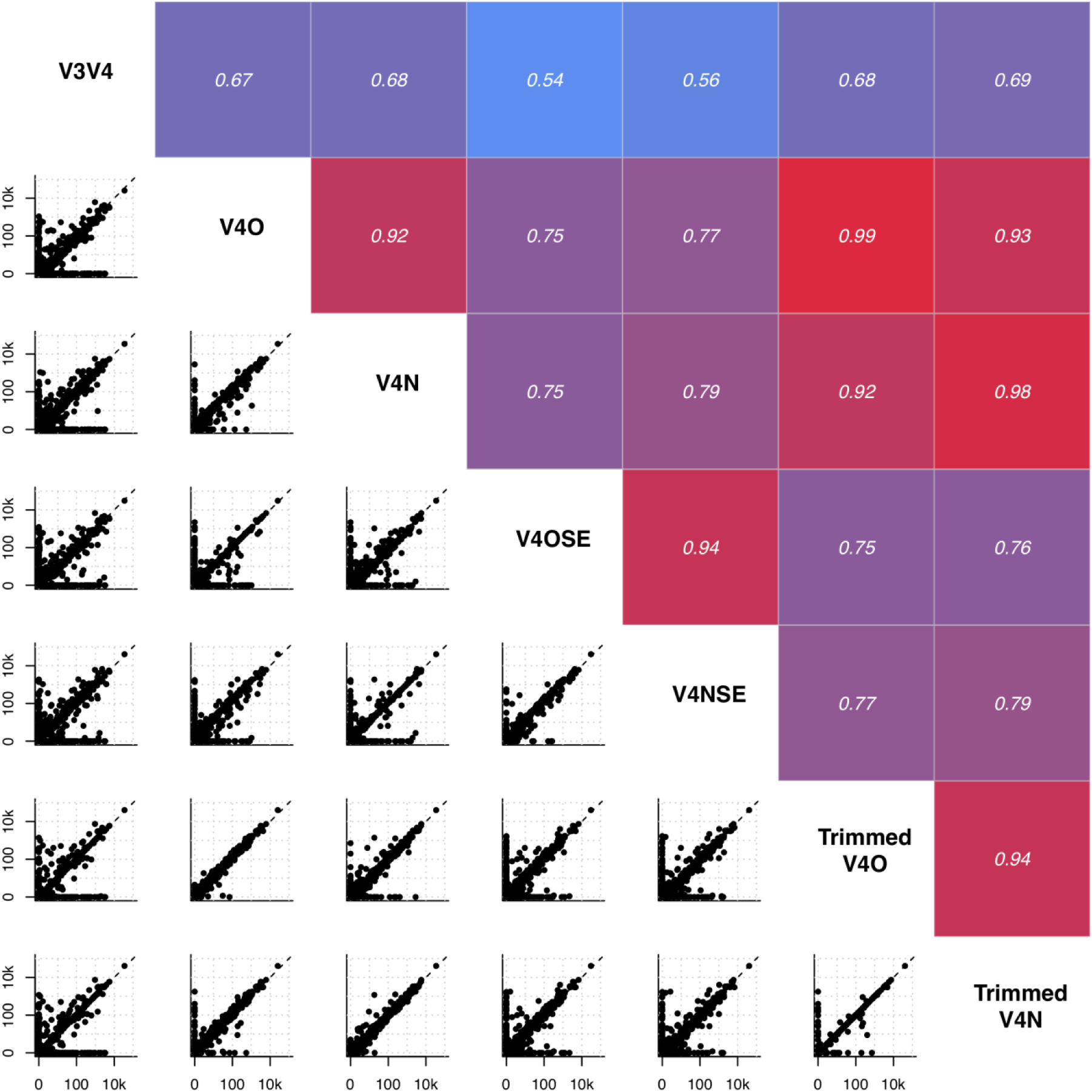
Taxonomic abundance correlations of seven analytical approaches. Lower left panels, pairwise taxonomic abundance correlations. Each point is an amplicon sequence variant shared by any two-analysis sets. Upper right panels, Pearson’s correlation coefficients for pairwise taxonomic abundance correlations; all tests were significant. Abbreviations are as in Fig. 3.

## DISCUSSION

The increasing popularity and advances in sequencing technology prompt human and clinical microbiome studies. However, the types of sequencing technology and reagents used limit meaningful comparisons of the obtained data. Here, we compared the most popular microbiome sequencing approaches that rely on the analysis of 16S amplicons (i.e., amplifying different hypervariable regions and coupling with DADA2 sequence variants’ denoising process). We evaluated the accuracy and consistency of 16S amplicons, which are captured by V3V4 and V4 regions. Our findings showed that V4O (515F-806R) primer yielded the most consistent, complementary, and accurate taxonomic profile. Additionally, we implemented an integrated approach for the V3V4 and V4 amplicons from different datasets by trimming V3V4 sequences up to the V4 region.

The NGS technology and HMP (HMP 1 and iHMP) generate massive sequencing data, promoting clinical microbiome association studies (33). These studies reveal numerous previously unknown host–microbe interactions that underlie various health issues, especially non-communicable diseases, with novel etiology linkages (3-7). For a population-based cohort study in the clinical microbiome research, data throughput and quality, the cost–benefit aspect, and validity of the analytical pipeline should all be considered. The 16S V3V4 and V4 hypervariable regions are widely selected for human microbiota profiling, but the fragments amplified from different regions should be coupled with suitable sequencing conditions.Accordingly, we here evaluated the compatibility of different amplicon libraries by *in silico* and empirical approaches, coupled with a denoising algorithm to detect ASVs. The analysis demonstrated that the current widely used amplicon primers capture over 80% of the taxonomic information. The amplicon sequence length was a critical factor for taxonomic profiling as longer sequences contain more genetic information than shorter sequences (11). The primers were another critical factor, as the amplicons are more representative of the sample complexity when generated by more universal (conserved) primers. For example, while the V3V4 and V4O approaches share nearly identical reverse primers, the V3V4 library was characterized by a higher ASV richness index (ASV numbers; Fig. S1A) but contained less taxonomic information than the V4O library (the archaeal phyla were almost absent in the V3V4 library; Fig. 2A and Table 3).

The taxonomic assignment accuracy is crucial for the subsequent study design, such as strain isolation, identification, and clinical or commercial applications (12). The *in silico* PCR analysis performed herein demonstrated that the V4O taxonomy assignment is more accurate than the V3V4-based assignments. However, even though the V4N-captured sequences overlapped with most regions captured by the V4O method, the taxonomic accuracy (at most 70% accuracy at phylum level) was much lower than that of either V3V4 or V4O method. In practice, the tV4O approach (V4O trimmed from V3V4) was compatible for integrating V3V4-generated data with V4O-generated data (Fig. 6) when testing the human gut microbiome samples, which contain fewer archaea than environmental samples. Since short-read amplicon sequencing only extracts partial information for the 1.5-kb 16S rRNA gene, the taxonomic identification is at most limited at approximately 80% accuracy at the genus level by V3V4 primer amplification and at the species level by V4O primer amplification.

The V4 amplicon sequencing meets economic benefits for microbiome studies. Amplicon sequencing in many studies, not only environmental ecological studies but also human-associated studies, focuses on the V4 region and relies on the EMP protocol. However, the library construction methods are restricted by the maximum read length of a sequencer (30). In other words, the amplified V4 amplicon fragment size (PCR product without the adapter-linking sequences) is expected to be approximately 291 bp if the library is constructed by following the Nextera XT two-step dual index PCR protocol. The two-step dual index PCR method was officially designed for 16S V3V4 library construction, coupled with MiSeq 600-cycle (300 PE) sequencing by Illumina (15). Although the two-step dual index PCR is not exclusive to MiSeq and V3V4 amplicons (34), its sequencing outputs (i.e., the read length and PE overlap) should be precisely calculated based on the quality score distributions from the 5’-end to the 3’-end of the read (35). For example, the PE reads overlap by less than 10 bp when a V4 amplicon library is constructed by using Nextera XT kit and sequenced as 150 PE. Most V4 150 PE reads do not pass the quality filtering and cannot be used to assemble both ends. In addition, the DADA2 algorithm only accepts PE reads with a more than 20-bp overlap as default (24).

Caporaso et al. (36) developed customized sequencing primers based on HiSeq 150 PE. The ensuing customized sequencing procedures entailed modification of the library construction methods (forward primers were directly linked to barcode sequences and an adapter), the sequencing program (using the TruSeq workflow and ignoring an error message from the sequencer system), and increasing the run by additional 14 sequencing cycles. Consequently, the method not only solved the unassembled read problem (yielding 253-bp amplicon fragments) but also lowered the sequencing cost per sample (Table 1). However, even though in some studies, the early stages of sequencing do not follow the EMP or Caporaso protocols, the unassembled reads can be analyzed by using a single-end pipeline. We here demonstrated that the single-end pipeline was comparable with the PE pipeline, with more than 90% confidence in the taxonomy assignment but a slightly inferior quantification (Pearson’s correlation coefficient 0.75) (Table 3).

The ASV denoising methods (23) (e.g., DADA2(24), Deblur(25), and UNOISE3(27)) are replacing the traditional operational taxonomic unit clustering methods for choosing representative amplicon sequences. These denoising methods detect real biological sequence features missed by clustering, and denoised features are specific and reproducible (23). Therefore, we suggest trimming the V3V4 and V4 sequences to the same region to acquire the same representative ASVs when combining different libraries. Although the V3V4 approach under-detected the archaea, the analysis of trimmed V4 amplicons recovered the qualitative and quantitative aspects of the human gut microbiome with over 90% confidence (Table 3 and Fig. 6).

In conclusion, in the current study, we evaluated the compatibility of 16S V3V4 and V4 amplicons typically analyzed in clinical microbiome profiling studies by using a sequence variant-denoising pipeline. Our findings suggest that: (1) the analysis of the PE V4O amplicon (amplified using primers 515F–806R) results in the most accurate taxonomic assignment; (2) the V3V4 amplicon analysis is compatible with the V4 amplicon analysis after trimming to the same region; and (3) while mid-high throughput sequencers reduce the cost of sequencing per sample, only a customized V4 library is suitable for stitching PE reads for subsequent analyses (36). The findings are empirical and analytical suggestions for cost-effective population-based or meta-analysis clinical microbiome studies.

## MATERIALS AND METHODS

### Ethics statement and sample collection

The studies involving human fecal sample collection and informed consent from human participants were approved by the Institutional Review Board of National Taiwan University Hospital, Taipei, Taiwan (201606045RINB). Fecal samples from 10 healthy volunteers were collected during February 2017 at the National Taiwan University, as described by Wu et al. (37).

### Primer selection and alternative V4 primer design

Two sets of 16S amplicon primers, which targeted the V3V4 (primers 341F–805R) and V4 (primers 515F–806R; V4O) regions, were selected from the Illumina-recommend (38) and EMP protocols (18), respectively. An alternative pair of V4 region-specific primers (V4N) consisted of modified primers of EMP V4 and Ghyselinck et al. (39): 519F, 5’-CAGCMGCCGCGGTAAT-3’, and 798R, 5’-GGGTWTCTAATCCKGTT-3’. The expected PCR product length was 279 bp. The V4N primers were synthesized with overhang adapters for index attachment and Illumina sequencing adapters(15). The coverage rate of each primer pair was evaluated by using *in silico* PCR simulation (see below).

### *In silico* PCR simulation

The read and taxonomy coverage rates of the SILVA 16S gene database (NR 132 99%) (31, 32) were evaluated by *in silico* PCR. The simulation was conducted by extracting the expected PCR fragments generated by amplification using the V3V4, V4O, and V4N primers. The *in silico* PCR pipeline was set-up using a UNIX shell script, as follows: (1) create a list of degenerate primer pairs, and link the forward primer to the reverse primer with “.*” from the 5’-end to the 3’-end (e.g., CCTACGGGAGGCAGCAG.*GGATTAGATACCCCAGTAGTC); (2) count the fragments extracted from the database fasta file by using UNIX command “*grep”* with the parameter *-c*; (3) obtain the targeted sequence fragments using the parameter *-o*; (4) extract the sequence ID using the parameter *-B 1*; (5) count the length of *in silico* PCR products using the *awk ‘{print length}’* UNIX command.

### Sequencing library preparation for mock community and human fecal microbiomes

#### DNA extraction

Genomic DNA from a mock community standard (ZymoBIOMICS Microbial Community Standard, catalog no. D6300, ZYMO RESEARCH, CA, USA) and from stool samples from 10 volunteers was extracted using QIAamp^®^ PowerFecal^®^ DNA Kit (QIAGEN, catalog no. 12830–50; Hilden, Germany). The genomic DNA was stored at m –20°C until PCR amplification and amplicon sequencing.

#### Amplification and NGS sequencing

Two-step PCR was performed, following the Illumina protocol for 16S metagenomic sequencing library preparation. PCR was first performed to capture the 16S V3V4 (primers 341F–805R) and V4 hypervariable regions (primers 515F– 806R and 519F–798R). Three libraries were then constructed by index PCR using the Nextera XT duel-Index PCR primers (15). The pooled libraries were PE-sequenced in the same run using the Illumina MiSeq reagent kit version 3 (San Diego, CA, USA) for 600 cycles at the Medical Microbiota Center of the First Core Laboratory, National Taiwan University College of Medicine.

### Bioinformatic analysis for microbial taxonomic profiling

#### Sequence denoising using DADA2 and the QIIME 2 pipeline

The sequences were processed by using QIIME 2 pipeline (version 2019.10) (9). The primer sequences were trimmed from the raw reads in the three libraries by using the *cutadapt* plugin. The trimmed single-end or PE sequences were subsequently denoised using the *DADA2* plugin in QIIME2. To obtain qualified ASVs, the reads were truncated from the 3’-end based on the quality score distribution to the following read length: (1) V3V4-forward, 270 bp, and V3V4-reverse, 210 bp; (2) V4O-forward, 131 bp, and V4O-reverse, 130 bp; and (3) V4N-forward, 134 bp, and V4N-reverse, 133 bp. In addition, theV3V4 reads were trimmed to the V4O and V4N read-length for further comparisons. High-confidence ASVs were then obtained by denoising, with quality filtering and chimera removal. The taxonomy was assigned using a naÏve Bayes classifier trained on the SILVA 132 99% full-length 16S rRNA gene sequence database (31, 32).

#### Microbial biodiversity and statistical analyses

All statistical analyses were conducted with R version 3.6.1(40). Microbial community analyses were performed using the *vegan* R package (41). Alpha diversity indices, including the Shannon index and Simpson index, were determined by using the “diversity” function; the Richness index was calculated by using the “specnumber” function. The beta diversity was determined based on the Bray–Curtis dissimilarity and visualized by principal coordinates analysis. ANOSIM was used to test the heterogeneity among individuals, controlling for the different primers. All univariate analyses were conducted by the Kruskal–Wallis test with α=0.05 cut-off for significance and Dunn’s test for post-hoc comparisons. Multiple-testing P-values were adjusted based on the false discovery rate by using the “p.adjust” function in R. The likelihood-ratio test (G-test with α=0.05 cut-off for significance) for the abundance profiles was performed by using the *RVAideMemoire* R package.

### Data availability

Sequences generated in the course of the current study have been deposited in the Sequence Read Archive (SRA) database under the accession number PRJNA643648.

## ACKNOWLEDGMENTS

We thank the research participants and research assistants from the Institute of Food Science and Technology, National Taiwan University, Taipei, Taiwan (Guan-Ling Ou) and National Taiwan University College of Medicine, Taipei, Taiwan (Yu-Tang Yang and Fang-Wei Kuo). We would also like to acknowledge the sequencing service provided by the Medical Microbiota Center of the First Core Laboratory, National Taiwan University College of Medicine, the computational resource support by Prof. Alex Hon-Tsen Yu at the Department of Life Science, National Taiwan University, and technical consulting by An-Chi Cheng at the University of Florida, Gainesville, FL, USA.

We declare that we have no competing interests.

P.Y.L. and W.K.W. conceived and planned the project. W.K.W., C.C.C., and S.P. were involved in sample collection, processing, and storage, and supervised the experiments. P.Y.L. conducted all bioinformatic and statistical analyses. P.Y.L. and W.K.W. were involved in data interpretation and manuscript planning. P.Y.L. drafted the manuscript. L.Y.S. and M.S.W. supervised the study. All authors approved submission of the final version.

## SUPPLEMENTARY FIGURE LEGENDS

### Supplementary Figures

**FIG S1.**
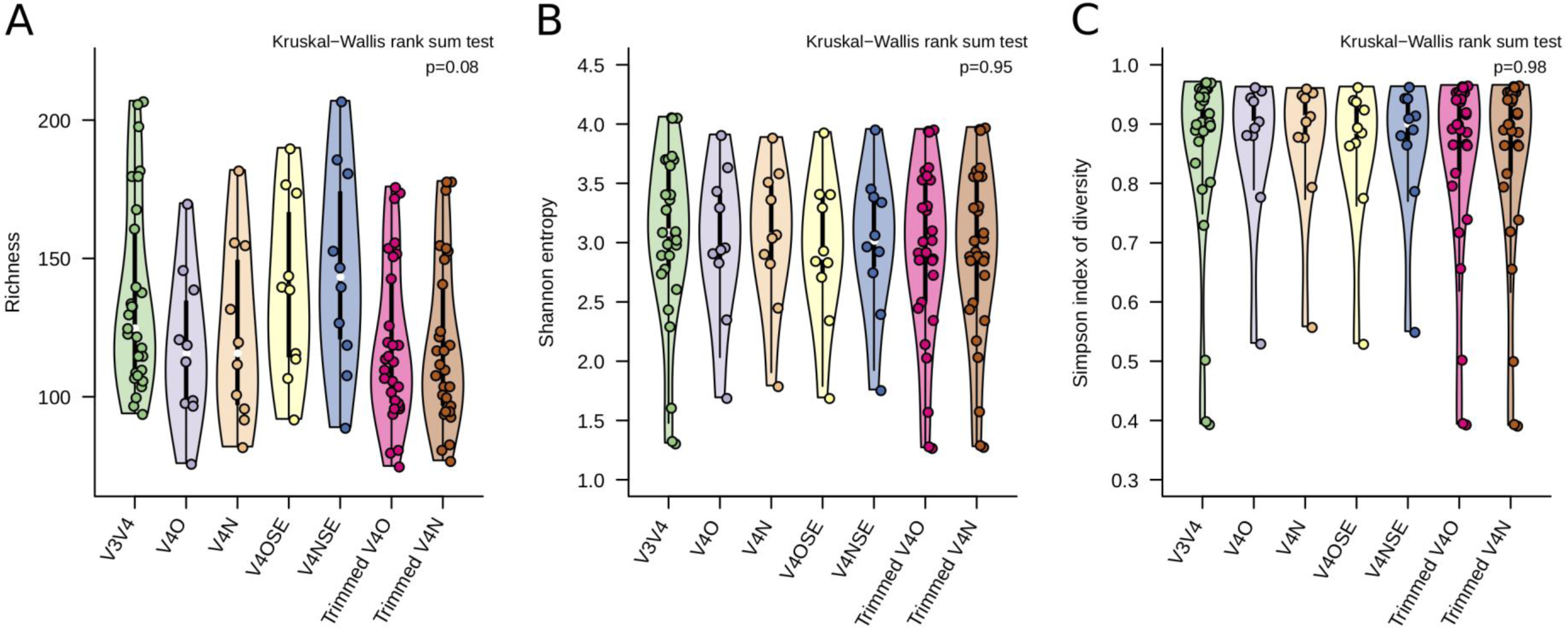
Alpha-diversity indices obtained with seven analytical approaches by testing the heterogeneity of 10 human fecal microbiomes. (A) Richness (amplicon sequence variant number). (B) Shannon entropy. (C) Simpson index of diversity. All indices were tested by Kruskal–Wallis rank sum test with alpha = 0.05. The seven analytical approaches were as follows: paired-end V3V4 amplicons (V3V4), paired-end V4O amplicons (V4O), paired-end V4N amplicons (V4N), single-end V4O amplicons (V4OSE), single-end V4N amplicons (V4NSE), and V4 amplicons trimmed from V3V4 (trimmed V4O, tV4O; and trimmed V4N, tV4N).

**FIG S2.**
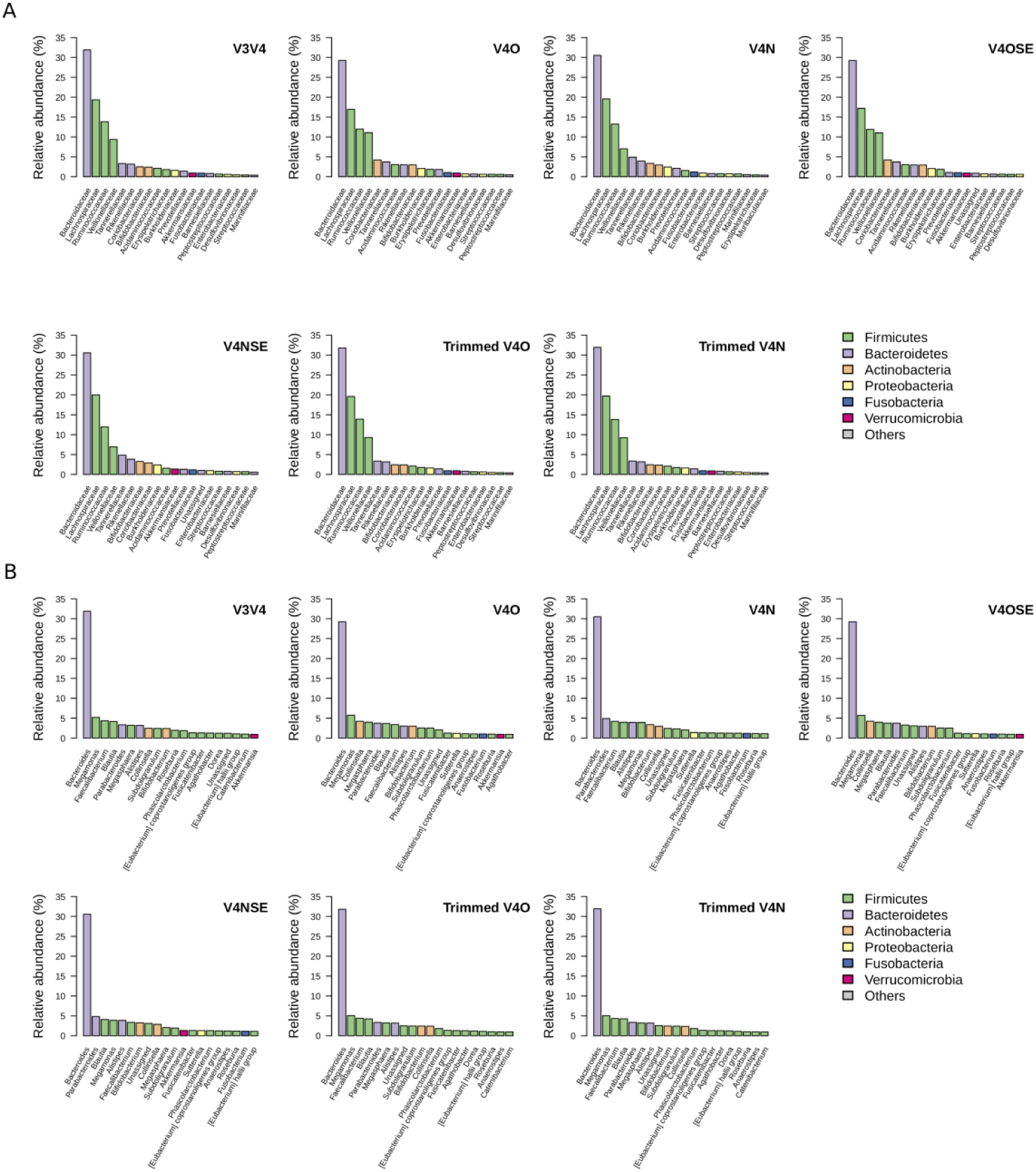
Taxonomic rank abundances obtained with seven analytical approaches by testing the heterogeneity of 10 human fecal microbiomes. (A) Family rank abundance. Bar fill color corresponds to the phylum. (B) Genus rank abundance. Abbreviations are as in Fig. S1.

### Supplementary Table

**TABLE S1.** *In-silico* archaeal sequence capture rates of the V3V4 and V4 primers

|  | **Number of archaeal sequences** | **Percentage of captured archaea within dataset** |
| --- | --- | --- |
| SILVA NR132<br>99 represented sequences | 19,501 | 5.27% |
| V3V4 | 69 | 0.024% |
| V4O | 15,916 | 5.26% |
| V4N | 8320 | 3.40% |
| tV4O | 65 | 0.024% |
| tV4N | 11 | 0.005% |

